# The maternal and fetal metabolic and immune landscapes of gestational diabetes mellitus

**DOI:** 10.1101/2024.04.13.589341

**Authors:** Duan Ni, Ralph Nanan

## Abstract

**Objectives:** Gestational diabetes mellitus (GDM) is the most common pregnancy-related medical complication. It is characterized by the development of hyperglycaemia during pregnancy and is known to lead to higher risk of metabolic disorders and other pathologies in both mothers and offsprings. Some studies probed the impacts of GDM, focusing on specific organs like placenta or adipose tissue, but so far, a systematic overview is lacking. Here, we aimed to curate a comprehensive atlas from currently available transcriptomic data for GDM, to comprehensively unravel how GDM influences the metabolic and immune landscapes in affected pregnancy.

**Methods:** RNA-sequencing (RNA-seq) data for maternal subcutaneous and omental fat, peripheral blood mononuclear cells (PBMCs), and fetal umbilical vein endothelial cells (HUVECs), amniocytes and cord blood mononuclear cells (CBMCs); and single-cell RNA sequencing (scRNA-seq) data for placenta and CBMCs were collated from previous publications. Comparative analyses and gene set enrichment analyses (GSEA) were carried out for the control versus GDM pregnancy.

**Results:** Maternal metabolic landscapes were consistently shifted by GDM, with reduced oxidative phosphorylation and fatty acid metabolism in maternal adipose tissues and PBMCs. GDM also caused inflammation solely in maternal subcutaneous fat. scRNA-seq analysis of placenta revealed that GDM reduced granulocytes and myelocytes but increased extravillous trophoblast cells. GDM also differentially impacted the metabolic and immune signals in different placental cell subsets. Contrarily, metabolisms in fetal compartments were minimally influenced by GDM. However, they consistently exhibited elevated inflammatory signals.

**Conclusion:** GDM differentially reprogrammed the maternal and fetal metabolisms and immunity.

---

Gestational diabetes mellitus (GDM) is the most common pregnancy-associated medical complication, not only increasing the risk of other pregnancy-related pathologies, but also predisposing both mother and offspring to metabolic disorders like diabetes, obesity and cardiovascular diseases (1, 2). GDM is characterized by abnormal gestational hyperglycaemia, and insulin-regulated metabolism. Previous studies have also found some GDM-induced changes in various maternal organs, including reprogrammed carbohydrate and fat metabolisms in placenta (3-5), and chronic inflammation in adipose tissue (6). The effect of GDM on the offspring is less well-described, apart from empirical observations on long-term metabolic conditions.

Moreover, most aforementioned findings are from stand-alone studies, a more systematic overview covering different compartments, and reconciling the maternal and fetal aspects, is lacking. Here, we surveyed currently available transcriptomic studies in GDM and collated datasets for comparative analyses for maternal compartments including adipose tissues (subcutaneous and omental fat) (7), peripheral blood mononuclear cells (PBMCs) (8) and placenta (9); and for fetal tissues including umbilical vein endothelial cells (HUVECs) (10), amniocytes (11) and cord blood mononuclear cells (CBMCs) (12) (Figure 1A, Table S1). We specifically profiled the metabolic and immune changes using gene set enrichment analysis (GSEA) (13), given their implications in GDM.

**Figure 1.**
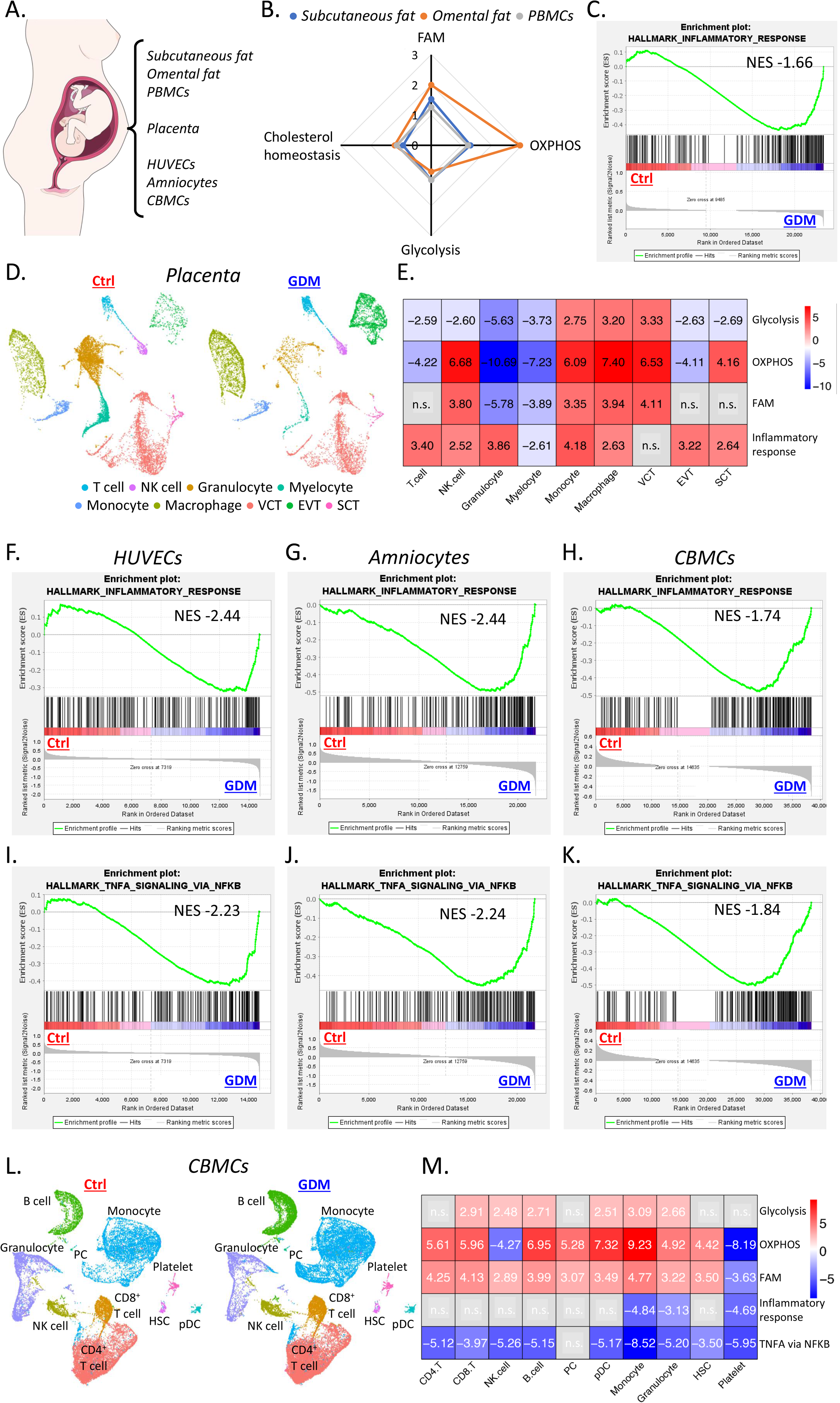
**A**. An overview of the tissue compartments covered in current study. **B**. Radar plot for the gene set enrichment analysis (GSEA) enrichment scores for fatty acid metabolism (FAM), oxidative phosphorylation (OXPHOS), glycolysis and cholesterol homeostasis gene sets in control compared to gestational diabetes mellitus (GDM) subcutaneous fat (blue), omental fat (orange) and peripheral blood mononuclear cells (PBMCs, grey). **C**. GSEA showing enrichment of inflammatory response gene set in GDM subcutaneous fat versus control. **D**. Uniform manifold approximation and projection (UMAP) plots visualizing the control (left) and GDM (right) placental cellular compositions analyzed by scRNA-seq. **E**. Heatmap visualizing the GSEA enrichment scores for glycolysis, OXPHOS, FAM and inflammatory response gene sets in different populations from control versus GDM placenta. Enrichment scores were specified, with positive values denoted enrichment in control while negative values denoted enrichment in GDM sample, and n.s. denoted no significance. **F-H**. GSEA showing enrichment of inflammatory response gene set in GDM human umbilical vein endothelial cells (HUVECs, **F**), amniocytes (**G**), and cord blood mononuclear cells (CBMCs, **H**). **I-K**. GSEA showing enrichment of TNFA_signaling_via_NFKB gene set in GDM HUVECs (**I**), amniocytes (**J**) and CBMCs (**K**). **L**. UMAP plots visualizing the control (left) and GDM (right) CBMC compositions analyzed by scRNA-seq. **M**. Heatmap visualizing the GSEA enrichment scores for glycolysis, OXPHOS, FAM, inflammatory response and TNFA_signaling_via_NFKB gene sets in different populations from control versus GDM CBMCs. Enrichment scores were specified, with positive values denoted enrichment in control while negative values denoted enrichment in GDM samples, and n.s. denoted no significance.

As shown in Figure 1B, relative to GDM, adipose tissues and PBMCs from healthy mothers consistently exhibited enhanced oxidative phosphorylation (OXPHOS) and fatty acid metabolism (FAM) signals, while other metabolism-related gene sets like glycolysis and cholesterol homeostasis were less affected (Table S2). One of the hallmarks of diabetes is the chronic inflammation. Similarly, inflammatory response gene set was enriched in GDM subcutaneous fat (Figure 1C), but interestingly not observed for omental fat. Together, these data revealed that GDM systematically shifted the maternal metabolic landscapes in multiple compartments, dampening OXPHOS and FAM, but triggered inflammation solely in subcutaneous fat.

Situating at the maternal and fetal interface, placenta is likely to play a critical role in GDM manifestation. Harnessing a previously published scRNA-seq dataset (9), we attempted to inspect at a single cell level how GDM affected different cell subsets within placenta, particularly their metabolic and immune profiles. scRNA-seq data was clustered and grouped into 9 main cell types including macrophages, monocytes, myelocytes, granulocytes, T cells, natural killer cells, villous cytotrophoblast cells (VCT), extravillous trophoblast cells (EVT) and syncytiotrophoblastcells (SCT) (Figure 1D). Similar to the original report (9), B cells were found in very low abundance. They were thus not included in further analysis, together with cells that could not be annotated to known cell types based on cell-type-specific marker genes (Others) (Figure S1). All 9 cell populations were present in both healthy and GDM placenta, but GDM was found to have higher proportion of EVT but lower granulocytes and myelocytes.

GSEA was run for different populations, comparing healthy control versus GDM patients (Figure 1E). GDM generally resulted in upregulation of glycolysis, except in monocytes, macrophages and VCT. It also promoted OXPHOS in T cells, granulocytes, myelocytes and EVT, but suppressed OXPHOS in others. FAM seemed to be less vulnerable to GDM and was increased in granulocytes and myelocytes but decreased in NK cells, monocytes, macrophages and VCT. Intriguingly, inflammatory signals were enriched in healthy control when compared to GDM samples for most populations, except for myelocytes and VCT. Collectively, the scRNA-seq analysis revealed that GDM conferred multifaceted impacts on placental cells. It altered their metabolic and immune profiles in patterns distinct to the aforementioned maternal compartments.

We next probed into the impacts of GDM on the fetus. In contrast to above maternal analyses, minimal difference was found for metabolic pathways comparing healthy and GDM samples by GSEA in HUVECs, amniocytes, and CBMCs. Instead, enrichments in inflammation-related gene sets in GDM were consistently detected (Figure 1F-H). This seems to be driven by tumor necrosis factor (TNF)-mediated responses, as the “TNFA_signaling_via_NFKB” gene set was enriched in all GDM fetal tissues (Figure 1I-K).

A published CBMC scRNA-seq dataset was next leveraged to gain more insights towards the impacts from GDM (12). As shown in Figure 1L, 10 cell subsets were identified in CBMCs, including CD4^+^ T cells, CD8^+^ T cells, B cells, plasma cells (PC), NK cells, monocytes, granulocytes, plasmacytoid dendritic cells (pDC), hematopoietic stem cells (HSC) and platelets. Similar to a prior report (12), population sizes of CBMC subsets were mildly affected.

scRNA-seq analysis casted more detailed insights towards the metabolic impacts conferred by GDM (Figure 1M). GDM dampened glycolysis in CD8^+^ T cells, NK cells, B cells, pDC, monocytes and granulocytes. For OXPHOS and FAM, they were generally higher in healthy controls in all subsets other than NK cells and platelets. scRNA-seq analysis supplied unprecedented insights towards the aforementioned elevated inflammatory signals during GDM. It unravelled a profound effect from the innate immune system. GDM significantly promoted the inflammatory response gene set in monocytes and granulocytes, the main immune constituents in CBMCs. Notably, platelets from GDM-affected CBMCs exhibited enhanced inflammatory signals as well. More strikingly, all CBMC populations except PC displayed elevated TNF-mediated signalling, consistent with the RNA-seq results. Together, these results highlighted that GDM conferred less pronounced effects on the metabolism of infants, but surprisingly promoted inflammatory signalling.

In this study, we casted unprecedented systematic insights towards the influences from GDM on both the mother and the offspring. GDM differentially modified the metabolism networks in the mother but to a less extent in the offspring. At the maternal-fetal interface, the placental metabolisms were rewired in a cell type-specific way, but its inflammatory signal seemed to be blunted. Notably, inflammatory pathways were activated in GDM-affected offsprings.

Here, we confirmed increased inflammation in maternal subcutaneous fat (6). This did not extend to omental fat, possibly hinting their differential physiological roles. Contrarily, inflammatory signals were consistently enhanced in GDM-impacted infants, possibly increasing their propensity to chronic inflammation, and consequently predispositions to inflammatory diseases (12, 14) and metabolic disorders (1, 2). It would be of great interests to interrogate whether these changes are due to epigenetic modifications and how long they persist over lifespan.

These findings seem to highlight the intermediate role of the placenta, separating the maternal and fetal distinct metabolic and immune landscapes. This potentially results in differential GDM-induced pathological sequels, requiring further investigations. Considering the importance and heterogeneity of the placenta, future spatial-scale analysis is warranted for more in-depth characterizations.

Finally, current offspring analyses substantially lack insights towards other metabolic and immune compartments, like the gastroenteric system. Here, materials like exfoliated intestinal epithelia cells might provide novel opportunities for mechanistic interrogation (15, 16).

Collectively, we presented a transcriptomic atlas interrogating the maternal and fetal metabolic and immune landscapes during GDM, supplying unprecedented insights for future related both basic and clinical research.

