## Supplementary Information for "The maternal and fetal metabolic and immune landscapes of gestational diabetes mellitus"

**Materials and Methods**

**Sex as a biological variable**

Data used in current study were collated from previous publications and their demographic background information was reported in the original papers (Table S1).

**Data curation and processing**

RNA-seq and single cell RNA-seq (scRNA-seq) data were curated from the previous publications as described in Table S1. RNA-seq data was normalized with DESeq2 (v1.34.0) (1). The resulting data was used for further analysis. scRNA-seq data was analyzed with *Seurat* (v4) (2, 3).

**Data analysis and statistics**

Gene set enrichment analysis (GSEA) was carried out with GSEA (v4.3.2) software (4) and *fgsea* package following their tutorials. scRNA-seq data analysis was carried out with the *Seurat* (v4) package (2, 3) following the analytical workflows in the original publications.

**Study approval**

N/A

**Data availability**

Data used in current study are all publicly available as described in Table S1.

**Supplementary Tables and Figure**

**Table S1.** Overview of the transcriptomic data used in current study.

| **Compartments** | **Analytical method** | **GEO ID** | **REF** |
| --- | --- | --- | --- |
| **Subcutaneous fat** | RNA-seq | GSE188799 | (5) |
| **Omental fat** | RNA-seq | GSE188799 | (5) |
| **Peripheral blood mononuclear cell** | RNA-seq | GSE92772 | (6) |
| **Placenta** | scRNA-seq | GSE173193 | (7) |
| **Umbilical vein endothelial cell** | RNA-seq | GSE49524 | (8) |
| **Amniocyte** | RNA-seq | GSE150621 | (9) |
| **Cord blood mononuclear cell** | scRNA-seq | GSE212309 | (10) |

**Table S2.** Gene set enrichment analysis (GSEA) scores for fatty acid metabolism, oxidative phosphorylation, glycolysis, cholesterol homeostasis, and inflammatory response gene sets comparing control and gestational diabetes mellitus (GDM) subcutaneous fat, omental fat and peripheral blood mononuclear cells (PBMCs). Positive values denote gene set enrichment in control samples while negative values denote gene set enrichment in GDM samples. n.s.: not significant.

|  | **Fatty acid metabolism** | **Oxidative phosphorylation** | **Glycolysis** | **Cholesterol homeostasis** | **Inflammatory response** |
| --- | --- | --- | --- | --- | --- |
| **Subcutaneous fat** | 1.58 | 1.32 | n.s. | n.s. | -1.66 |
| **Omental fat** | 2.00 | 2.95 | n.s. | 1.21 | n.s. |
| **PBMCs** | 1.27 | 1.23 | n.s. | n.s. | n.s. |

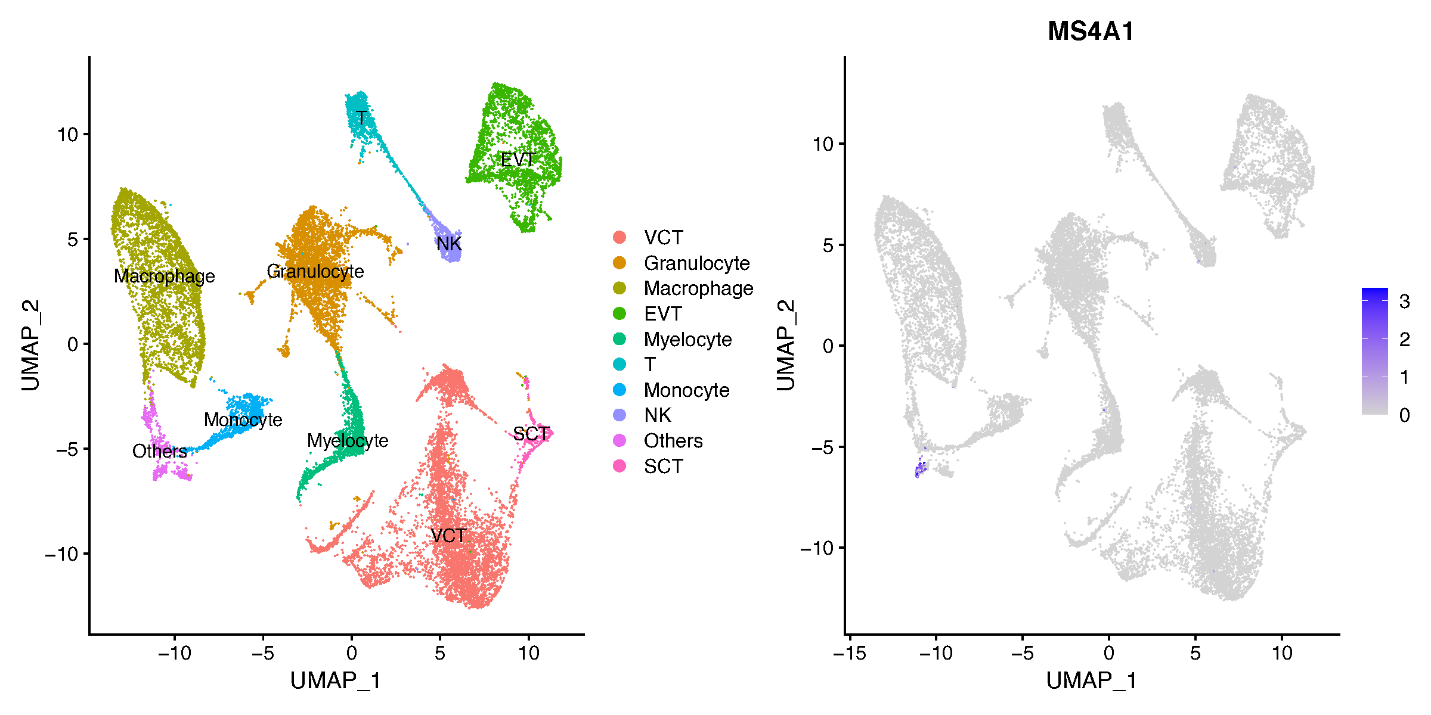

**Figure S1.** Uniform manifold approximation and projection (UMAP) plot visualizing the cellular compositions of control and gestational diabetes mellitus (GDM) placenta analyzed by scRNA-seq (left) and feature plot for the expression of B cell marker *MS4A1* (right).
